# Apoptosis in snowflake yeast: novel trait, or side effect of toxic waste?

**DOI:** 10.1101/029918

**Authors:** Jennifer T. Pentz, Bradford P. Taylor, William C. Ratcliff

## Abstract

Recent experiments evolving de novo multicellularity in yeast have found that large-cluster forming genotypes also exhibit higher rates of programmed cell death (apoptosis). This was previously interpreted as the evolution of a simple form of cellular division of labor: apoptosis results in the scission of cell-cell connections, allowing snowflake yeast to produce proportionally smaller, faster-growing propagules. Through spatial simulations, Duran-Nebreda and Solé (2015) develop the novel null hypothesis that apoptosis is not an adaptation, per se, but is instead caused by the accumulation of toxic metabolites in large clusters. Here we test this hypothesis by synthetically creating unicellular derivatives of snowflake yeast through functional complementation with the ancestral ACE2 allele. We find that multicellular snowflake yeast with elevated apoptosis exhibit a similar rate of apoptosis when cultured as single cells. We also show that larger snowflake yeast clusters tend to contain a greater fraction of older, senescent cells, which may explain why larger clusters of a given genotype are more apoptotic. Our results show that apoptosis is not caused by side effects of spatial structure, such as starvation or waste product accumulation, and are consistent with the hypothesis that elevated apoptosis is a trait which co-evolves with large cluster size.

## Introduction

Duran-Nebreda and Solé (2015) recently reported results from a computational model examining the evolution of multicellularity in experimentally-evolved ‘snowflake’ yeast. In this paper, they offer an explanation for a pair of experimental results from Ratcliff et al. (2012): First, some yeast cells within clonal clusters undergo programmed cell death (apoptosis). Early snowflake yeast are small and exhibit little cell death, but sustained directional selection for faster settling resulted in the evolution of both greater cluster size and proportionally higher rates of apoptosis (Ratcliff et al., 2012). Second, the rates of apoptosis observed within snowflake yeast clusters increased nonlinearly with cluster size. Duran-Nebreda and Solé are able to recapitulate these observed dynamics in a simulation model in which apoptosis is induced by the build-up of waste products, such as acetate. All else being equal, they hypothesize that larger clusters experience greater diffusional gradients, causing internal cells to be exposed to sufficient quantities of waste products to induce cellular suicide. Elevated apoptosis in large clusters is thus, in their view, caused by a pre-existing genotype by environment interaction that emerges once large cluster size evolves (Duran-Nebreda & Solé, 2015).

This intriguing hypothesis contrasts with the hypothesis put forward by Ratcliff et al. (2012), which posits that apoptosis in large-bodied snowflake yeast is an evolved trait with a genetic basis. Specifically, Ratcliff et al. (2012) suggest that apoptosis is an adaptation to large cluster size that helps mitigate slower growth rates, which is caused by poor resource diffusion to internal cells. Apoptosis severs cell-cell connections, resulting in branch scission and the production of a propagules (Ratcliff et al., 2015). Production of proportionally smaller propagules is adaptive as long as their faster growth more than compensates for reduced survival at the end of the 24 h culture cycle. Indeed, evolutionary simulations show that this criteria is often met, as elevated apoptosis readily evolves *in silico* (Conlin & Ratcliff, In press; Libby et al., 2014). The distinction between these hypotheses is of crucial importance for properly interpreting Ratcliff et al. (2012)’s experimental results: if elevated apoptosis is an adaptation, not a side-effect, it demonstrates that selection can readily favor the evolution of altruistic cellular traits in simple clusters of cells. Such selection underpins the evolution of cellular division of labor (Bourke, 2011; Fisher et al., 2013; Hammerschmidt et al., 2014; Libby & Rainey, 2013; Michod et al., 2006; Ratcliff et al., 2012; Shelton & Michod, 2014; Smith & Szathmáry, 1995), and is thus of fundamental importance for understanding the evolution of increased multicellular complexity.

Early (isolated after 7 days of settling selection) and late (from 60 days of selection) snowflake yeast were obtained from the experiment conducted in Ratcliff et al. (2012). These yeast gained the ability to form clusters through the inactivation of the trans-acting transcription factor *ACE2* (Ratcliff et al., 2015), which regulates the expression of a number of enzymes involved in mother-daughter cell separation after mitosis (King & Butler, 1998; Oud et al., 2013; Voth et al., 2005). As a result, daughter cells remain attached to mother cells, creating the branched ‘snowflake’ phenotype. The 7-day yeast form relatively small clusters with little apoptosis, while the 60-day yeast form large clusters with elevated apoptosis (Ratcliff et al., 2012). To test Duran-Nebreda and Solé’s hypothesis that stronger diffusional gradients in larger clusters is the proximate cause of increased apoptosis, we conducted an experiment to examine apoptosis in the absence of any multicellular structure. We created unicellular derivatives of the 7- and 60-day snowflake yeast by complementing the evolved *ace2* strains with a functional copy of the ancestral *ACE2* allele. The resulting unicellular strains differ only at a single locus (*ACE2*) from their snowflake ancestors, but do not experience an accumulation of waste products due to multicellular structure. 60-day snowflake yeast retained their elevated apoptosis even when grown as single cells, demonstrating that apoptosis is not simply a side-effect of diffusional limitation in large clusters. To explain the nonlinear increase in apoptosis and cluster size observed *among clonal clusters* of the 60-day snowflake yeast strain (Ratcliff et al., 2012, Figure 5A), we simulate snowflake yeast growth and reproduction, and show that larger clusters tend to develop a core of old, potentially apoptosis-prone cells, while small propagules are typically composed of young and healthy cells. We find that older yeast are disproportionately apoptotic, supporting our hypothesis that large clusters contain more apoptotic cells because they contain more aged, senescent cells, not due to the accumulation of toxic metabolites.

## Methods

*Creating unicellular yeast from snowflake genotypes.* We obtained unicellular derivatives of isolates from early (7 days of settling selection) and late (60 days of settling selection) snowflake yeast from the experiment conducted in Ratcliff et al. (2015). Briefly, we replaced a single copy of the naturally-evolved nonfunctional *ace2* in each isolate with the ancestral *ACE2* allele fused with the antibiotic resistance gene *KANMX4* (Ratcliff et al., 2015) using the LiAc/SS-DNA/PEG method of yeast transformation (Gietz et al., 1995). Transformants were then plated on solid Yeast Peptone Dextrose medium (YPD; per liter: 20 g dextrose, 20 g peptone, 10 g yeast extract and 15 g agar) with 200 mg/L of the antibiotic G418. For each transformant, the insertion location of the transformation sequence was confirmed by PCR.

*Measuring apoptosis in yeast.* We measured the rate of apoptosis in early and late isolates of multicellular snowflake yeast as well as their respective unicellular derivatives, along with unicellular controls evolved without selection for faster settling. These latter strains were grown under the same conditions as snowflake yeast (daily 1:100 dilution into 10 mL YPD, 250 RPM shaking at 30°C), but did not experience any settling selection. To measure apoptosis, we developed a novel method to count the fraction of apoptotic cells directly. Uni- and multicellular yeast were dual-labeled with the green-fluorescent reactive oxygen species stain dihydrorhodamine 123 (DHR 123; 2.5 mg/mL in 95% ethanol) as well as a blue-fluorescent vacuole stain, CellTracker™ Blue CMAC, allowing direct measurement of each (Figure 1B and C). Before measuring apoptosis, all strains were struck out on solid YPD from -80°C stocks, then grown in 10 mL liquid YPD for 24 h at 30°C, 250 RPM shaking. Ten replicate populations of each yeast strain were then initiated with a 1:100 dilution into 10 mL YPD then cultured for 12 h. After 12 h of growth, yeast were stained with a 1% solution of DHR 123 (Madeo *et al*., 1999) and a 0.1% solution of CellTracker™ Blue CMAC, and incubated in the dark for two hours. Yeast were then double-washed with carbon free yeast nitrogen base (YNB) buffer (Sigma Aldrich, 6.7 g/L) and unicellular strains diluted 10-fold, while multicellular strains were diluted two-fold. 5 μL of this culture was placed between a slide and a 25 × 25 mm coverslip. Placing such a small volume of media on the slide tends to flatten multicellular clusters (Figure 1A), helping keep all cells in focus (Figures 1B and C). Large-field mosaic images were collected for each isolate by combining 36 separate images (each collected at 100 × magnification) using a Nikon Eclipse Ti inverted microscope with a computer-controlled Prior stage, resulting in a 10185 × 7665 pixel composite image. This technique results in large sample sizes, is sensitive to dim fluorescence, and minimizes sample bias arising from single images in which yeast touching the edge of the field of view are discarded (this disproportionately affects large clusters). Counts of all cells (CellTracker™ Blue CMAC; green dot labels in 1B) and apoptotic cells (DHR 123; blue dot labels in 1C) were acquired using the spot detection algorithm in the NIS-Elements v. 4.30 software package.

**Figure 1.**
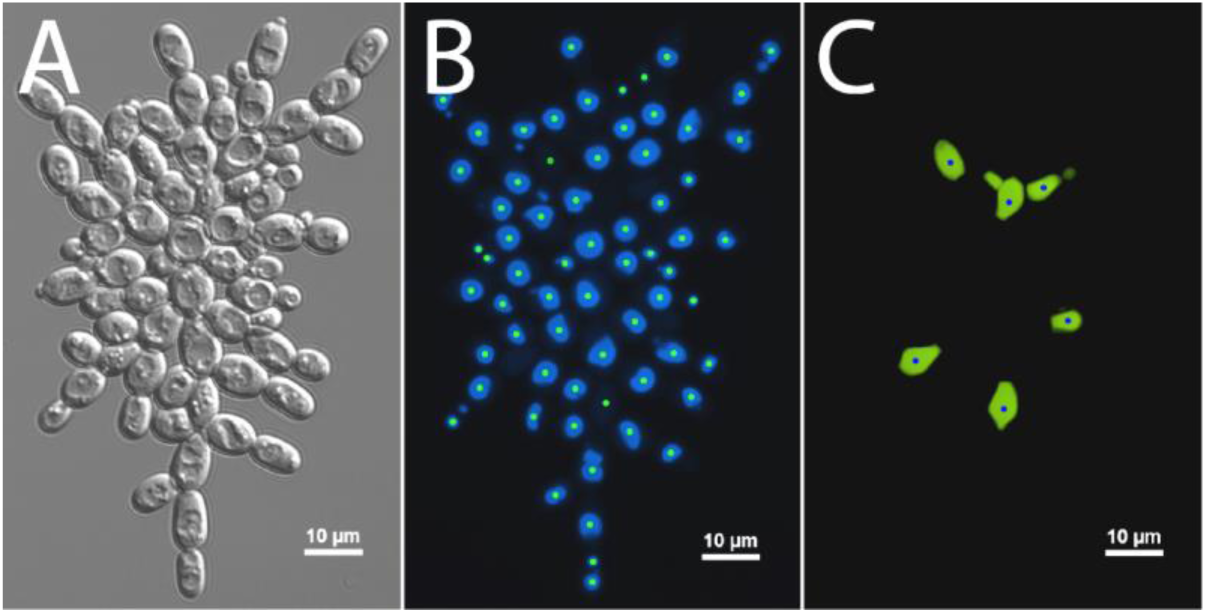
Measuring apoptosis in snowflake yeast. Pictured is the same snowflake cluster imaged with A) Differential Interference Contrast microscopy, B) the blue fluorescent vacuole stain CellTracker™ Blue CMAC, and C) the green fluorescent apoptosis stain dihydrorhodamine 123. We obtained counts of all cells by counting the number of vacuoles (green dots marking cells in B) and apoptotic cells (blue dots marking cells in C) by using the spot detection algorithm in NIS-Elements v. 4.30. The same method was used to measure apoptosis in unicellular strains.

*Measuring the age of apoptotic yeast.* We measured the age of non-apoptotic and apoptotic cells by dual-labeling unicellular derivatives of early and late snowflake yeast with DHR 123 (to determine which cells are undergoing apoptosis) and the chitin-binding stain calcoflour (1 mg/mL in water). Calcoflour labels “bud scars” left on the mother cell after mitotic division, allowing us to determine the replicative age of the cell (Mortimer & Johnston, 1959). Unicellular derivatives of both 7- and 60-day snowflake yeast were grown and stained with DHR 123 as described above. After the 2 h incubation to label apoptotic cells, yeast were stained with a 1% solution of calcoflour and incubated in the dark for an additional 5 minutes. Yeast were then double-washed with YNB and diluted 10-fold. 5 μL was then placed between a slide and a 25 × 25 mm coverslip and imaged at 400 × magnification. 6 μm thick z-stack images of 100 apoptotic and 100 non-apoptotic cells were made by imaging 6 slices with a 1 μm step size. Bud scar counting was performed on the maximum intensity projections. To improve the efficiency of our counts, we processed these images using a custom Python script that subtracted background fluorescence and increased the contrast of bud scar edges.

*Graph-based simulation.* Cellular growth and scission were stochastically simulated using the Gillespie algorithm (Gillespie, 1977). Given a current state of the system, represented by a population of graphs, the algorithm proceeds by randomly sampling the time the next event occurs, Δ*t*, and by randomly sampling which event occurs (cell reproduction via node growth or propagule production via edge scission). Because we do not have a precise understanding of how snowflake yeast clusters reproduce, we modeled branch scission with two separate approaches. With weighted reproduction, edge scission depends on the amount of strain associated with that node, or its weight *w_i_*, calculated as the total number of downstream path-connected cells (*e.g*., total number of genealogical descendants connected to each cell; see edge values in Figure 3A). This weighting parameter increases linearly with the number of attached cells, so scission tends to occur towards the center of the cluster. In contrast, with unweighted separation, scission occurs at a constant rate throughout the cluster, so edges tend to separate close to the cluster periphery where most nodes are located. Cell growth events occur at a constant rate, *g_j_*=1, and edge removal events occur either at a constant rate or at a rate proportional to the current weight, *r(w_i_).* The time the next event occurs is obtained by sampling from an exponential probability density function (PDF), *PDF* (Δ*t*) *= λe*^−λΔ^*^t^*, parametrized by the total sum of rates across nodes and edges in the population, where:

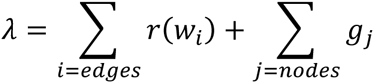

The probability of choosing an event is the ratio of its rate to the total rate. In the unweighted model, edges were removed at a constant rate, *r_i_* = ***ϕ***, while in the weighted model edges were removed in a weight (*w_i_*)-dependent rate, *r_i_(w_i_)* = ***ρ*** *w_i_.*

## Results and Discussion

Growth form (unicellular vs. multicellular) did not directly affect apoptosis. Early (7 day) snowflake yeast exhibited similar low rates of apoptosis (0.48% and 0.45%) when grown as snowflakes or unicells, respectively (Figure 2B). The 60-day snowflake yeast strain evolved a four-fold increase in apoptosis (1.90%), an elevated rate that was maintained in the unicellular form (1.61%). As a control, we examined apoptosis in a unicellular yeast transferred for either 7 or 60 days without settling selection, and which remained unicellular. This control maintained its low ancestral rate of apoptosis (0.30% vs. 0.34% for day 7 and day 60 strain, respectively). Differences in apoptosis were assessed via a full factorial two-way ANOVA, with days of evolution and treatment as independent variables and percent apoptosis as the dependent variable (*F*_5,54_=92.56, *p*<0.0001, *r*^2^=0.9). Between-group differences were assessed with Tukey’s HSD; significance at α=0.05 is denoted by different letters (a or b) in Figure 2B. Overall, only 60-day snowflake yeast possessed greater rates of apoptosis, and this did not depend on whether they were grown as snowflake clusters or as single cells. These results demonstrate that the elevated rates of apoptosis observed in late snowflake yeast isolates cannot be attributed to diffusional gradients imposed by multicellularity, and supports the hypothesis that apoptosis is a trait co-evolving with increased cluster size.

**Figure 2.**
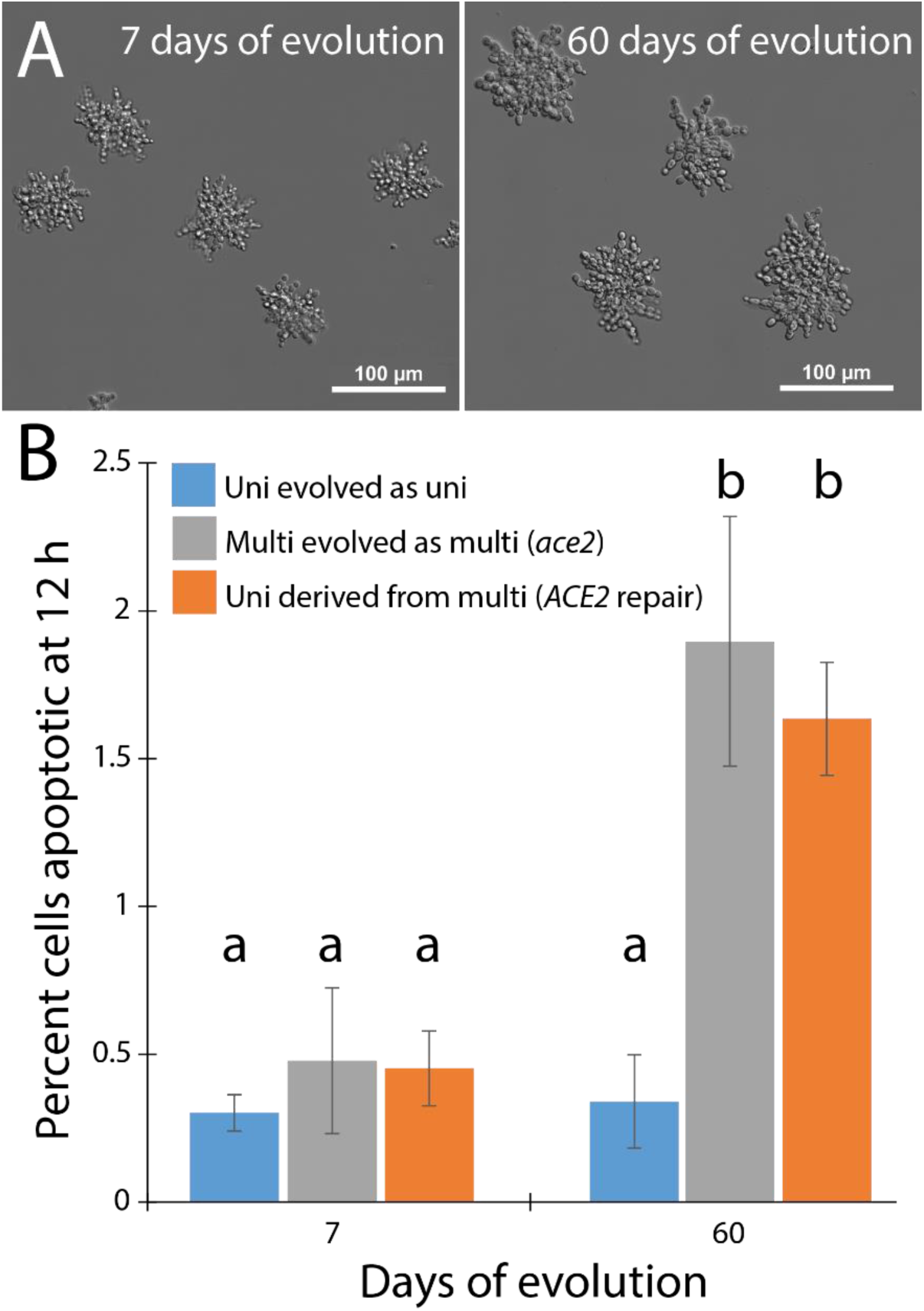
Elevated apoptosis is not a consequence of cluster spatial structure. A) Snowflake yeast from the same population at different time points. Cluster size increased between 7 and 60 days. B) 60-day snowflake yeast show a four-fold increase in rate of apoptosis over 7-day snowflake yeast (grey bars; 1.90% vs. 0.48% apoptotic, respectively). Unicellular derivatives (transformed with a functional *ACE2* allele) maintained this elevated rate of apoptosis (orange bars; 1.61%). A control line in which we evolved unicellular yeast without selection for faster settling (it stayed unicellular) maintained its low ancestral rate of apoptosis (0.34% vs. 0.30% for 60- and 7-day isolates, respectively). Shown are means ± the standard deviation of 10 biological replicates.

If apoptosis is not caused by waste product accumulation, then how do we explain the nonlinear increase in apoptosis seen among larger clusters of a single genotype (Figure 5a of Ratcliff et al. 2012)? We hypothesize that elevated apoptosis is due to the accumulation of old, senescent cells in large clusters, which itself is due to the geometry of snowflake yeast and their mode of reproduction. Snowflake yeast clusters reproduce by branch scission. Liberated branches act as propagules, which grow in size until they are large enough to produce their own offspring (Ratcliff et al., 2012). One effect of this mode of reproduction is that propagules tend to contain many young, recently-produced cells. In contrast, the cells in the interior of the cluster are less likely to be released by branch scission, and thus may stay in the cluster center for considerably longer than those on the periphery. We modeled this effect quantitatively. The structure of a snowflake yeast cluster can be represented as a graph with a tree-like structure, in which cells are nodes and cell-cell connections are edges (Figure 3A; Libby et al., 2014; Ratcliff et al., 2015). Clusters reproduce when an edge is severed. We considered 512-cell snowflake yeast clusters (9 generations of growth from a single cell), and calculated the fraction of cells older than four generations in both propagules and parent clusters with varying reproductive asymmetry (Figure 3B). Parent clusters always contained a larger fraction of old cells; an effect that was enhanced by greater reproductive asymmetry. Apoptosis is a well-known consequence of aging in yeast (Herker et al., 2004; Laun et al., 2001), so the difference in mean age of cells in propagules relative to large, mature clusters may explain the higher rates of apoptosis observed in the latter.

**Figure 3.**
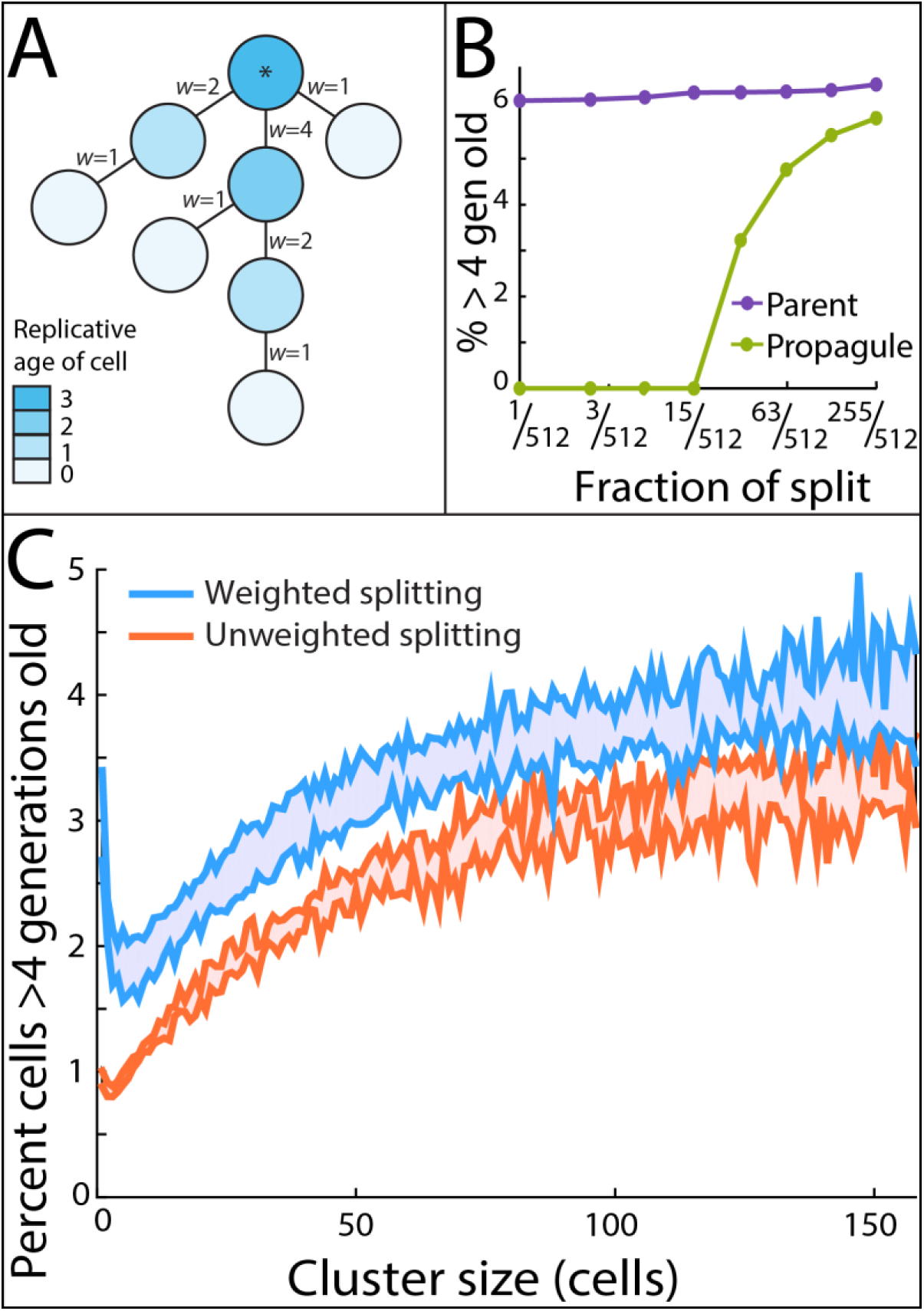
Larger clusters contain a greater fraction of old cells. A) A snowflake yeast cluster represented by a graph of nodes and edges. Nodes represent component cells in the cluster and edges represent physical connections between a pair of cells. The root cell (denoted by *) is the oldest cell in the cluster, with younger cells being closer to the periphery. Edge weights *w* depend on the number of attached cells. B) Larger propagules tend to contain a higher proportion of old cells. This is due to the geometric structure of snowflake yeast: propagules that separate from the parent cluster contain mostly young cells, skewing the age distribution of old clusters and increasing the proportion of old cells they contain. C) We simulated a population ofsnowflake yeast growing for nine divisions with two models: in the first the per-edge probability of scission depends linearly on the number of attached descendent cells (weighted splitting; see *w* values in A), while in the second it is position independent (unweighted splitting). This figure shows only common size classes, (*i.e*., those that occurred in at least 10% or the simulations). For both weighted and unweighted splitting, we plot the minimum and maximum percentage of cells older than 4 generations obtained in all simulations (10^3^ runs for each parameter, ***ϕ*** and ***ρ***).

We next simulated the growth and reproduction of snowflake yeast over a period of time corresponding to nine cellular generations and examined the relationship between cluster size and cellular age. We simulated our dynamic model across a range of values for ***ϕ*** [.04 .0425 .045 .0475 .05] and ***ρ*** [.01 .0125 .015 .0175 .02], with 10^3^ simulations per parameter value. Our results confirm a positive relationship between cluster size and proportion of older cells (Figure 3C); these results were similar with both the weight dependent and independent models of edge scission. Indeed, the ~2% increase in cells over 4 generations old between small and large clusters (Figure 3C) is similar to the ~2% difference in apoptosis seen in small vs. large snowflake yeast clusters of the 60-day strain (Ratcliff et al., 2012, Figure 5A). The increase in old-aged single cells in the weighted model (Figure 3C) is due to a statistical anomaly: edge scission is rare in large clusters for peripheral cells with *w*=1, as they have low weight, so many singletons were the result of a split in an early round of growth when clusters contained just a few cells. Because cellular reproduction is stochastic, many singletons at the end of the simulation are old cells which failed to reproduce.

To determine if apoptosis is an age-dependent process in our 60-day snowflake yeast, we measured the replicative age of 100 apoptotic and 100 non-apoptotic cells by counting bud scars. Older cells were increasingly apoptotic (Figure 4; *p*=0.0217, *r*^2^=0.61, linear regression). To summarize, we found that snowflake yeast that evolved elevated rates of apoptosis after 60 days of selection continue to display this trait even when genetically altered to grow as single cells, disproving the hypothesis that side effects of cluster spatial structure (*e.g.,* waste product accumulation or starvation) directly cause this phenotype. We hypothesize that the positive within-genotype relationship between cluster size and fraction of cells undergoing apoptosis reported in Ratcliff et al. (2012) is due instead to differences in the age of the cells within these clusters. Larger clusters tend to contain a higher percentage of old cells than smaller clusters, and old cells are more likely to undergo apoptosis.

**Figure 4.**
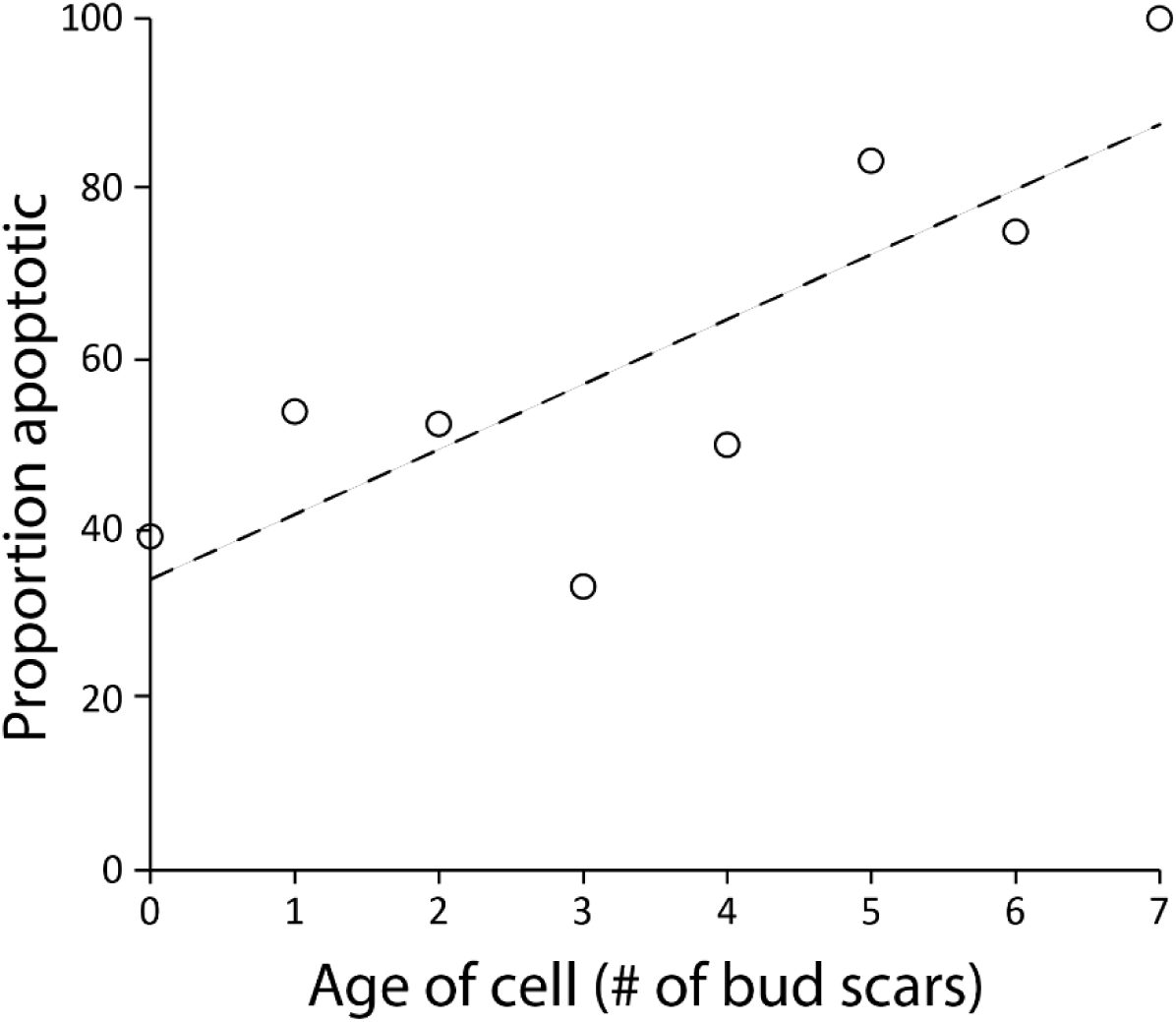
Old-aged cells are disproportionately apoptotic. We measured the age of 100 apoptotic and 100 non-apoptotic cells. Old cells were more likely to be apoptotic than young cells (y = 7.6x + 34.2, *p*=0.02, *r*^2^=0.61, linear regression).

While our experimental results contradict Duran-Nebreda and Solé’s (2015) main conclusions, their work remains relevant and important. Indeed, their physical model established a plausible null hypothesis for elevated apoptosis and has deepened our understanding of the snowflake yeast experimental system. More generally, nascent multicellular organisms are expected to be heavily affected by physical affects arising from their spatial structure (Drasdo et al., 2007; Libby et al., 2014; Newman & Bhat, 2008; Ratcliff et al., 2015), and physical models will continue play a critical role in the study of this major evolutionary transition.

## Author contributions

J.P. and W.C.R. designed the experiments and analyzed the data. B.T. wrote the model. W.C.R., J.P., and B.T. wrote the paper. The authors have no conflicts of interest to declare.

## Acknowledgements

The authors would like to thank Joshua Weitz for helpful feedback on the manuscript. This work was supported by NASA Exobiology Award #NNX15AR33G.

